# Persistent cell proliferation signals correlates with increased glycolysis in tumor hypoxia microenvironment across cancer types

**DOI:** 10.1101/2020.03.16.993311

**Authors:** Jinfen Wei, Kaitang Huang, Meiling Hu, Zixi Chen, Yunmeng Bai, Shudai Lin, Hongli Du

**Author notes:** These authors contributed equally (Jinfen Wei, Kaitang Huang). Correspondence address should be addressed to Hongli Du.

## Abstract

**Background:** Altered metabolism is a hallmark of cancer and glycolysis is one of the important factors promoting tumor development. Given that the absence of multi-sample big data research about glycolysis, the molecular mechanisms involved in glycolysis or the relationships between glycolysis and tumor microenvironment are not fully studied. Thus, a more comprehensive approach in a pan-cancer landscape may be needed.

**Methods:** Here, we develop a computational pipeline to study multi-omics molecular features defining glycolysis activity and identify molecular alterations that correlate with glycolysis. We apply a 22-gene expression signature to define the glycolysis activity landscape and verify the robustness using clinically defined glycolysis samples from several previous studies. Based on gene expression signature, we classify about 5552 of 9229 tumor samples into glycolysis score-high and score-low groups across 25 cancer types from The Cancer Genome Atlas (TCGA) and demonstrate their prognostic associations. Moreover, using genomes and transcriptome data, we characterize the association of copy-number aberrations (CNAs), somatic single-nucleotide variants (SNVs) and hypoxia signature with glycolysis activity.

**Findings:** Gene set variation analysis (GSVA) score by gene set expression was verified robustly to represent glycolytic activity and highly glycolytic tumors presented a poor overall survival in some cancer types. Then, we identified various types of molecular features promoting tumor cell proliferation were associated with glycolysis activity. Our study showed that TCA cycle and respiration electron transport were active in glycolysis-high tumors, indicating glycolysis was not a symptom of impaired oxidative metabolism. The glycolytic score significantly correlated with hypoxia score across all cancer types. Glycolysis score was also associated with elevated genomic instability. In all tumor types, high glycolysis tumors exhibited characteristic driver genes altered by CNAs identified multiple oncogenes and tumor suppressors. We observed widespread glycolysis-associated dysregulation of mRNA across cancers and screened out HSPA8 and P4HA1 as the potential modulating factor to glycolysis. Besides, the expression of genes encoding glycolytic enzymes positively correlated with genes in cell cycle.

**Interpretation:** This is the first study to identify gene expression signatures that reflect glycolysis activity, which can be easily applied to large numbers of patient samples. Our analysis establishes a computational framework for characterizing glycolysis activity using gene expression data and defines correlation of glycolysis with the hypoxia microenvironment, tumor cell cycle and proliferation at a pan-cancer landscape. The findings suggest that the mechanisms whereby hypoxia influence glycolysis are likely multifactorial. Our finding is significant not just in demonstrating definition value for glycolysis but also in providing a comprehensive molecular-level understanding of glycolysis and suggesting a framework to guide combination therapy that may block the glycolysis pathway to control tumor growth in hypoxia microenvironment.

## 1. Introduction

Altered metabolism is a hallmark of cancer^[1, 2]^. Rapidly proliferating tumor cells consume glucose at a higher rate compared to normal cells and part of their glucose carbon is converted into lactate, this is referred to as the ‘aerobic glycolysis’^[3]^ which has been correlated with advanced tumor progression^[4]^, treatment resistance^[5, 6]^ and poor clinical outcome^[7, 8]^ in various cancers. Numerous studies have confirmed that glycolysis intermediate can rapidly meet the energy requirement for cell proliferation, the material needs for fatty acids and nucleotides ^[9]^. The glycolysis products, lactic acid also plays multiple roles in cancer processes, including energy regulation, immune tolerance, wound healing, cancer growth and metastasis^[10, 11]^. These studies have established that glycolysis can provide favorable conditions for tumor proliferation and thus plays a pivotal role in cancer development.

Previous studies have showed that glycolysis effect seems to be a consequence to oncogenic signaling activation such as P53^[12]^, MYC^[13]^ and PI3K-AKT signaling^[14]^. However, the role of the microenvironment in driving metabolism alteration is increasingly recognized in recent years^[15, 16]^. Hypoxia is one of the key physiological and micro environmental differences between tumor and normal tissues. It induces DNA amplification^[16, 17]^ and increases clonal selection^[18]^, resulting in aggressive cancer phenotypes. A crucial results underlying such condition in a highly hypoxia tumor microenvironment is metabolic reprogramming of tumor cells^[19]^, such as, glycolysis is an adaptation to this low oxygen pressure microenvironment for activation by hypoxia-inducible factor HIF1A which can stimulate glycolytic by transactivation genes involved in SLC2A1 and ALDOA^[20]^. These prior studies provided interesting insights into the interplay between specific glycolysis genes with hypoxia environment or other molecular oncogenic signal. However, the main mechanisms driving the glycolysis shift remain unknown nor how glycolysis affects tumor progression remains incompletely understood. Therefore, there is an urgent need for understanding the comprehensive adjustment manner of regulation of glycolysis.

Although glycolysis is a targetable prognostic feature and there are already drugs targeted to key enzymes, the disappointing results of glycolysis-targeted therapy trials were observed in some cancer types ^[21, 22]^. Researchers have employed positron emission tomography (PET) following injection of the glucose analogue 18 F-fluorodeoxy-glucose (FDG) to diagnose tumour glycolytic ability and attempted to research the related mechanism^[23-25]^. However, not all cancers avidly take up FDG. Breast cancers, for example, show up to 20-fold differences in their FDG-PET signal that was attributed by histopathologic heterogeneity ^[26]^. Besides, FDG-PET cannot possible easily applied to large cohorts of patient samples with large data volume. Thus, there is still a lack of simple molecular characterization of glycolysis and comprehensive study of molecules related to tumor glycolysis. To fill this gap, we defined the glycolysis level using 22-gene expression and evaluated in 9,229 samples representing 25 distinct tumor types to create a pan-cancer quantification and explore glycolysis-associated molecular signatures in great depth. This is the first study to identify gene expression signatures that reflect glycolysis activity, which can be easily applied to large numbers of patient samples. We calculated the glycolysis distribution, compared the difference of clinical prognosis, genomic instability and key pathways between glycolysis-high and low tumors, and discovered the correlation between hypoxia and glycolysis across multiple tumor types. Our study strongly suggests that glycolysis promotes tumor proliferation, is influenced by a strong hypoxia pressure for specific molecular aberrations. Furthermore, it provides the theoretical basis for understanding tumor evolution in hypoxia microenvironment and suggests a framework to guide combination therapy that may block the glycolysis pathway to control tumor growth.

## 2. Result

### 2.1 Classification of tumor glycolytic activity by a gene expression signature and high glycolysis correlate with poor clinical features

To classify the glycolysis activity of tumor samples, we choose a 22-gene expression signature shown to represent the glycolytic activity and tested its robustness(see Methods). The samples with top 30% score was defined as glycolysis high tumors, bottom 30% was glycolysis low tumors according to glycolysis score by GSVA analysis (Fig.1A). In the 25 cancer types surveyed, both glycolysis score-high and glycolysis score-low groups contained ≥;30 samples (Supplementary Tab.1). To examine whether glycolysis score was driven by modest differences in the levels of many members or more dramatic effects on only 1 to 2 key enzymes, we determined the contribution of individual enzymes to overall pathway enrichment based on 22-gene expression in all tumor samples. Genes encoding enzymes showed the most consistent enrichment within the glycolysis score in pan-cancer (Supplementary Fig.1A) and also in one of the cancer types UCEC with the large sample size more than 500 (Fig.1B). We next tested whether 22 gene set expression could serve an “FDG signature” and predict glycolysis activity. We then performed analyses to validate its performance and assess the robustness of our signature using the GEO data in three independent gene expression datasets of cancer cell lines and tumor fragments with high and low glycolysis capacity determined by PET using 18 F-FDG uptake. We found a strong association between measured FDG uptake and glycolysis score (Fig.1C). The observed consistency suggests that glycolysis score based on 22-gene signature is appropriate to represent glycolysis activity in different cancer types.

**Fig.1:**
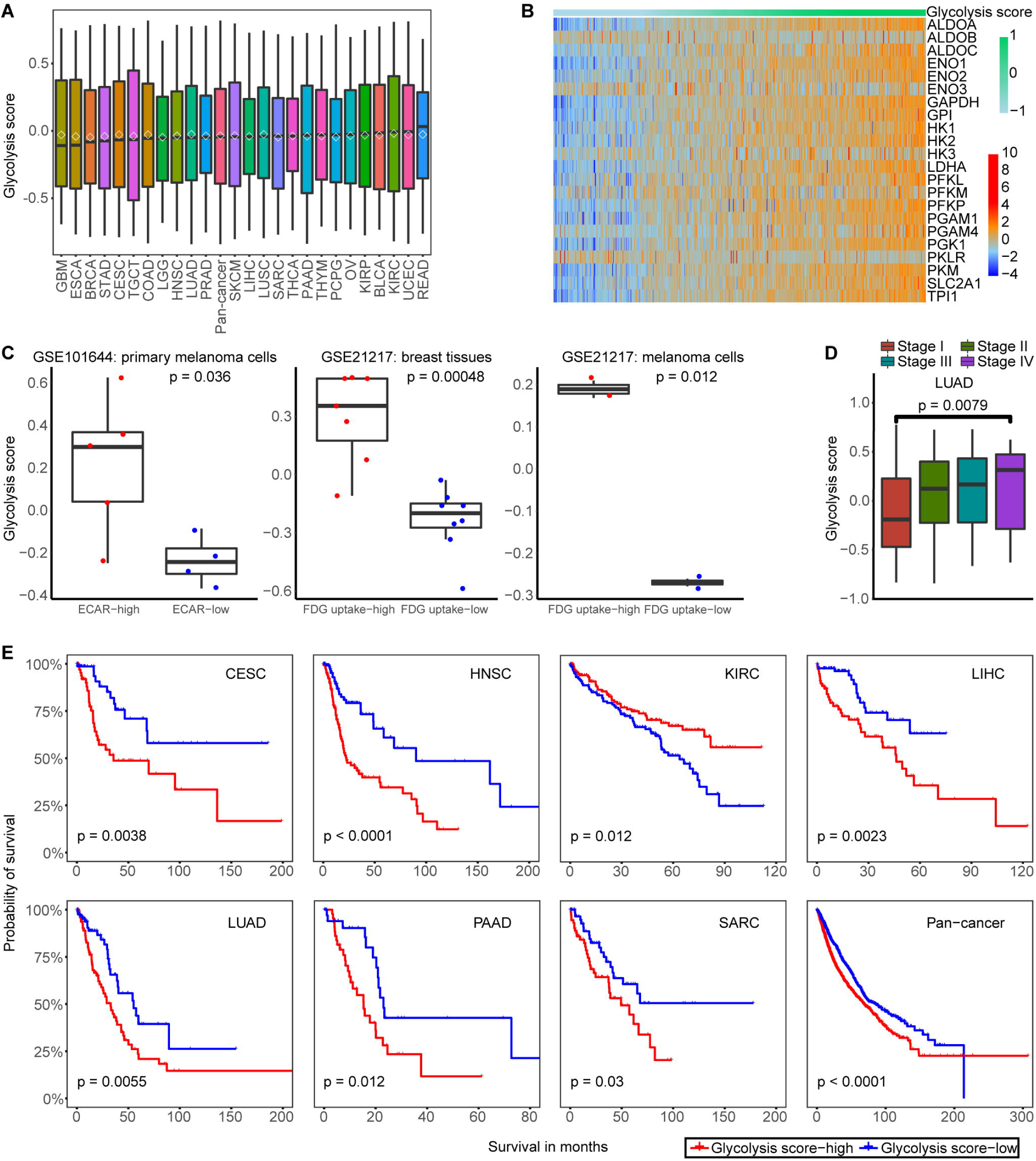
Validation of a 22-gene expression signature for glycolysis activity and clinical significance of glycolysis. A. Glycolysis scores based on the mRNA abundance signature for 25 tumor types, sorted by the median GSVA score (horizontal black line) and mean score (diamond pattern) for each cancer type. B. Samples are ordered from lowest to highest glycolysis score with 22-gene expression (z score of log2(TPM+1)) distribution in UCEC. The top color bar shows glycolysis score. C. Glycolysis scores of cancer cell lines and tumor fragments under FDG high uptake and low conditions in three datasets. A two-sided Student’s t-test was used to assess the difference. P□<□0.05. D. The glycolysis score was higher in clinical stage IV compared with stage I in LUAD, T-test was used to assess the difference. P□<□0.05. E. Kaplan–Meier curves show that glycolysis score-high status is associated with worse survival time in multiple cancer types. A two-sided log-rank test P□<□0.05 is considered as a statistically significant difference.

To examine whether cancers with the high and low glycolysis tumors were clinically distinct, we next determined glycolysis activity whether having correlations with patient overall survival, since survival represents a key clinical index of tumor aggressiveness. The higher glycolysis activity was associated with tumor advanced-stage in LUAD and BRCA (Fig.1D, Supplementary Fig.1B). We observed that glycolysis score-high tumors were consistently associated with lower overall survival in several cancer types in Kaplan Meier survival analysis with log-rank test, such as PAAD, LUAD and pan-cancer pattern (P < 0.05)(Fig.1E). These results highlighted the clinical relevance of metabolic subtypes in some cancer type and suggested the potential prognostic power of glycolysis activity classification.

### 2.2 The genomic alteration in glycolytic score high tumors

We next sought to identify genomic changes that characterize tumor glycolysis. To assess whether glycolysis was associated simply with an elevated CNV rate and SNVs features, we focused on three tumor types with large sample more than 500 (BRCA, LUAD and UCEC) (Fig.2A-C). In BRCA, high glycolysis tumors were more likely to harbor loss of REXO1 (adj. P < 10^−5^) and gain of MYC and ARSG (adj. P < 10^−10^) (Fig.2A). High glycolysis breast tumors also showed an elevated rate of TP53 point mutations (adj. P < 10^−10^) and reduced CDH1 mutations (adj. P < 10^−5^). Gain of LGI4 and ZBTB32 and loss of CDKN2A were observed in LUAD with high glycolysis (adj. P < 10^−2^) (Fig.2B). High glycolysis tumors in UCEC were associated with MUC16 and TTN mutations (adj. P < 10^−2^) (Fig.2C). Besides, alterations in several other genes were also associated with glycolysis in BRCA, LUAD and UCEC (Supplementary Tab.2).

**Fig.2:**
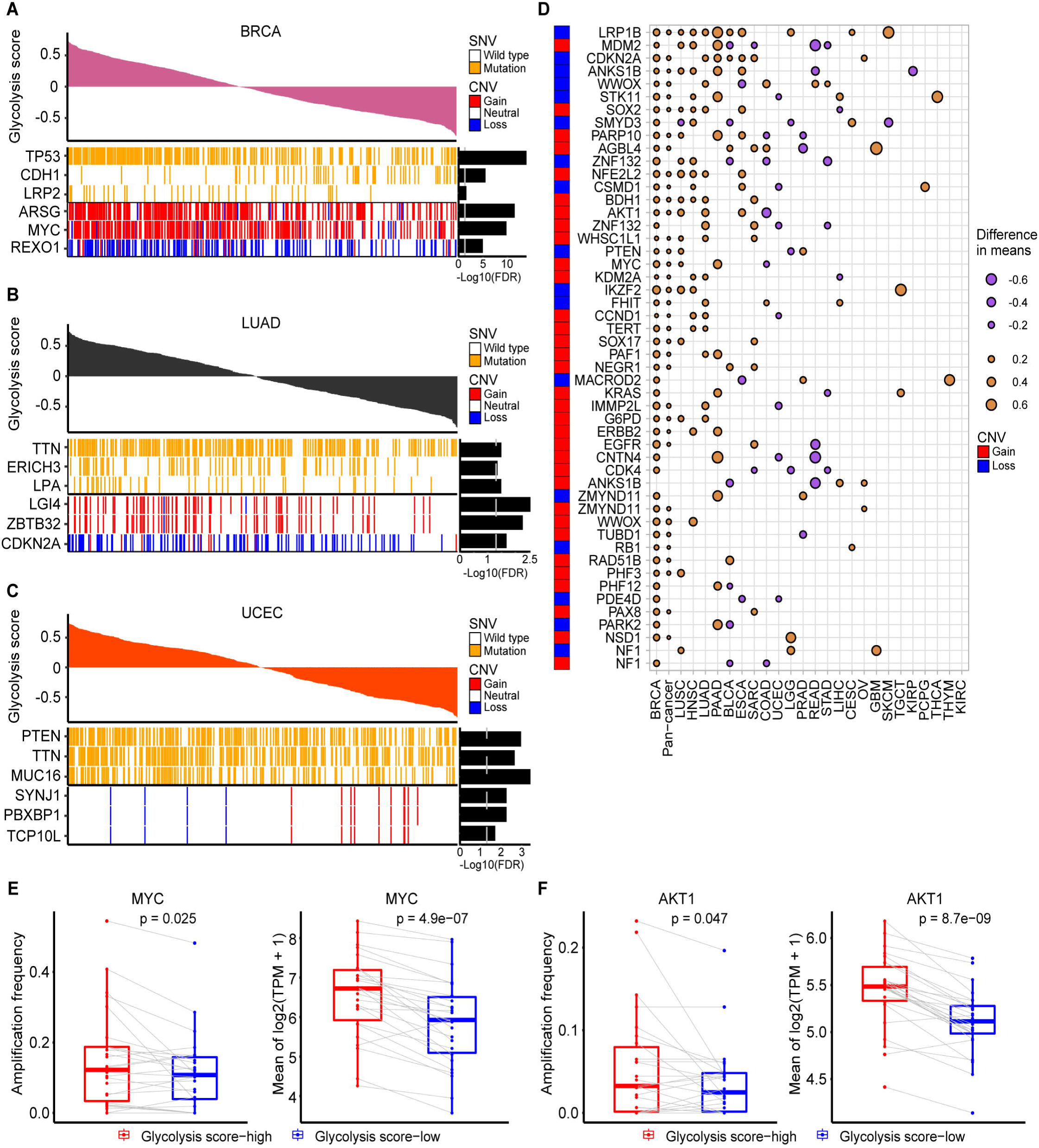
Characteristics of genomic changes associated with glycolysis activity. A-C. Notable associations of SNVs and CNAs with tumor glycolysis in subjects with BRCA (A), LUAD (B) and UCEC (C). Benjamini and Hochberg adj.P values are on the right (T test). D. Association of glycolysis with CNAs in oncogenes and tumor suppressor genes (T test). Dot size indicates the difference in mean glycolysis score between tumors with a CNA (gain for oncogene, loss for tumor suppressor gene) and those without a CNA. E. Amplification frequency of MYC and mean expression of MYC is higher in glycolysis-high than low tumors across 25 cancer types. F. Amplification frequency of AKT1 and mean expression of AKT1 is higher in glycolysis-high than low tumors across 25 cancer types. A paired Student’s t-test P□<□0.05 is considered as a statistically significant difference.

Next, analysis of 114 cancer driver genes altered by CNAs^[27]^ identified multiple oncogenes and tumor suppressors recurrently associated with glycolysis activity in 25 cancer types (Fig.2D, Supplementary Tab.3). As showed in Figure 2D, loss of the tumor-suppressor gene LRP1B was associated with elevated glycolysis in 11 separate tumor types, whereas gain of the AKT1 oncogene was associated with elevated glycolysis in five tumor types. In addition to BRCA, gain of MYC was also observed in PAAD and LUSC (P < 10^−2^). To explore how CNVs may influence these genes mRNA abundance, we compared the expression of these genes between glycolysis high and low tumors. We observed MYC and AKT1 mRNA expression was consistent with copy number amplification across cancer types (Fig.2E-F, Supplementary Fig.2A). Besides, correlation analysis showed MYC, AKT1 mRNA expression and glycolysis score were positive correlated across a variety of cancers ranging from r = 0.10 to 0.57 for MYC and r = 0.15 to 0.50 for AKT1 (P < 0.05) (Supplementary Fig.2B). In addition, tumor mutation load burden (TMB) was also associated with glycolysis activity in our analysis. High glycolysis samples had significantly increased TMB in 11 cancer types, such as BRCA, STAD and UCEC (Supplementary Fig.2C).

### 2.3 Cancer hallmarks, metabolic reprogramming and glycolysis activity

To explore the differences in metabolic characteristics and cancer hallmarks of the high and low glycolysis groups in each cancer type (Supplementary Tab.4), we did differential signature enrichment based on GSVA analysis, using publicly available gene set in Molecular Signatures Database (MSigDB) database. Using independently transcriptomic signatures, we sought after differentially enriched processes in the glycolysis high and low tumors (adj.P < 0.05 difference between the absolute means of GSVA scores in the two groups). The heat map combined with GSVA score difference showed that tumor proliferation signature, DNA replication, G2M checkpoint and MYC targets gene signatures were more active in high glycolysis groups across 23∼24 cancer types (Fig.3A). As expected, tumor cell proliferation signature was positively correlated with glycolysis activity in 23 cancer types(r = 0.11∼0.67)(Fig.3B, Supplementary Fig.3), implying that the high glycolysis tumor is characterized by a high proliferation rate. In addition, we observed a statistically significantly activation in cellular response to hypoxia signatures in 23 cancer types(Fig.3A). In the metabolism pathways, TCA cycle was significantly up regulated in glycolysis high groups in 20 cancer types. Besides, metabolism of nucleotides, nucleotide salvage, pentose phosphate pathway, glucose metabolism, nucleobase biosynthesis gene sets were enriched in glycolysis high groups across 24∼25 cancer types. Glutamate and glutamine metabolism, glycogen metabolism, mitochondrial fatty acid beta-oxidation and respiratory electron transport, metabolism of amino acids and derivatives and amino acids-serine biosynthesis were in glycolysis-high samples across 19-20 cancer types but not in THYM and GBM (Fig.3C).

**Fig.3:**
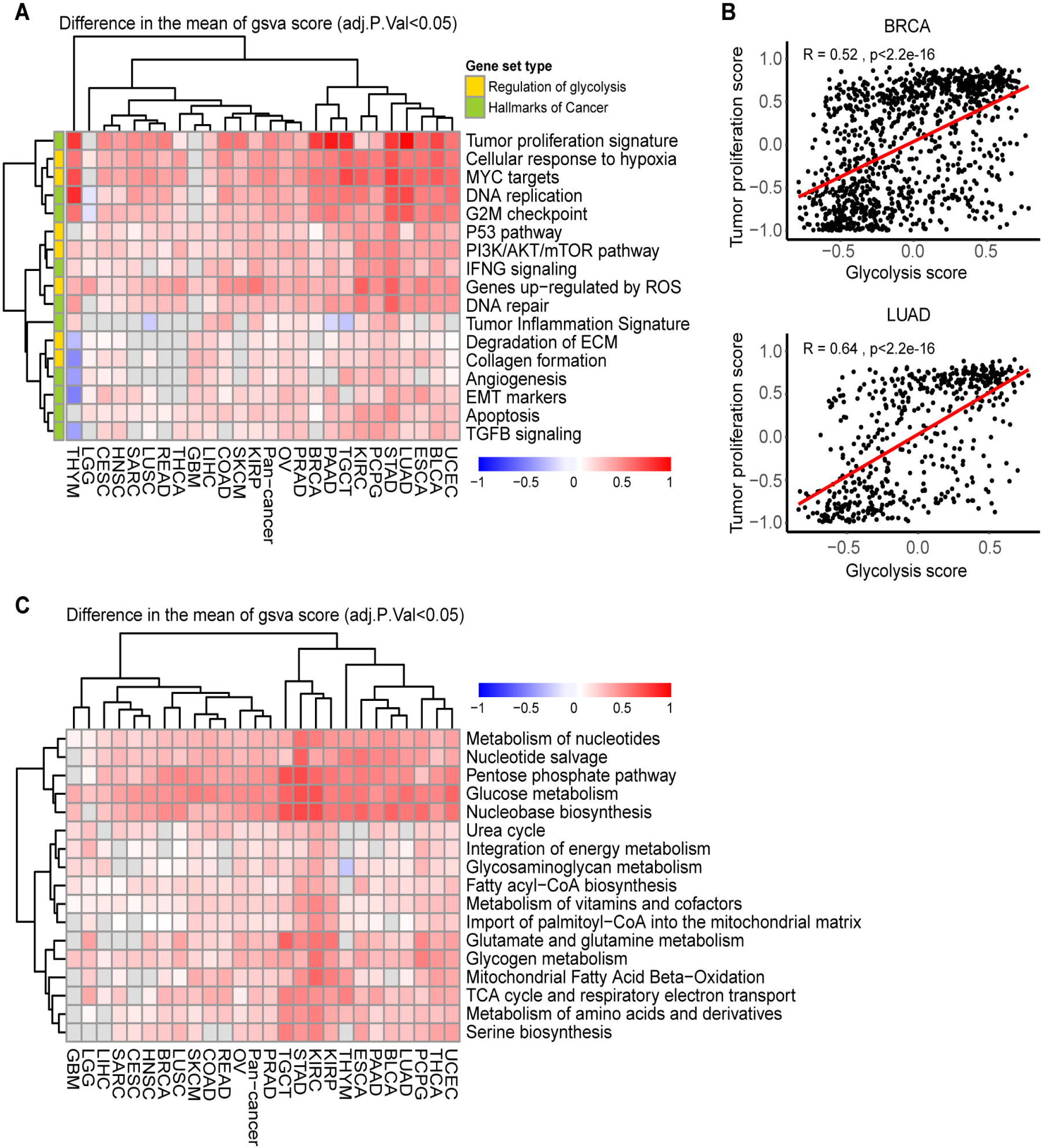
Association of glycolysis score with cancer hallmarks and metabolic reprogramming. A. Heatmap showing the difference of GSVA scores of cancer hallmarks signatures enriched in the high glycolysis versus low tumors. adj.P < 0.05. B. The spearman correlation between tumor proliferation signatures and glycolysis score in BRCA and LUAD, P□<□0.05. C. Heatmap showing the difference of GSVA scores of cancer metabolic reprogramming signatures enriched in the high glycolysis versus low tumors. adj.P < 0.05.

### 2.4 The association between glycolysis and hypoxia

Based on cellular response to hypoxia signatures was significantly active in glycolysis-high tumors shown in the above result (Fig.3A), we sought to determine whether there is a relationship between them through big data research. Since direct measurements of environment status of the cells are not available, the hypoxia status was defined by an established gene expression signature and was widely applied in previous researches^[28-30]^. GSVA analysis was also used to calculate the hypoxia score across cancer types. We compared hypoxia score in our glycolysis high and low groups, and found that hypoxia score was significantly higher in glycolysis-high tumors than the low one (Fig.4A, P < 2e-16). Next, by the calculation of spearman correlation, we found that glycolysis and hypoxia score were highly correlated with r > 0.7 across 25 cancer types (Fig.4B, Supplementary Fig.4). To exclude the possibility that the positive correlation was driven by a few correlated genes with high variation in expression levels while most other genes were not correlated, we did further correlation calculation between each gene in glycolysis set and hypoxia score or gene in hypoxia set and glycolysis score. The results confirmed that most of individual genes defining glycolysis score or hypoxia score were also positively associated with two scores (Supplementary Fig.5A-B).

**Fig.4:**
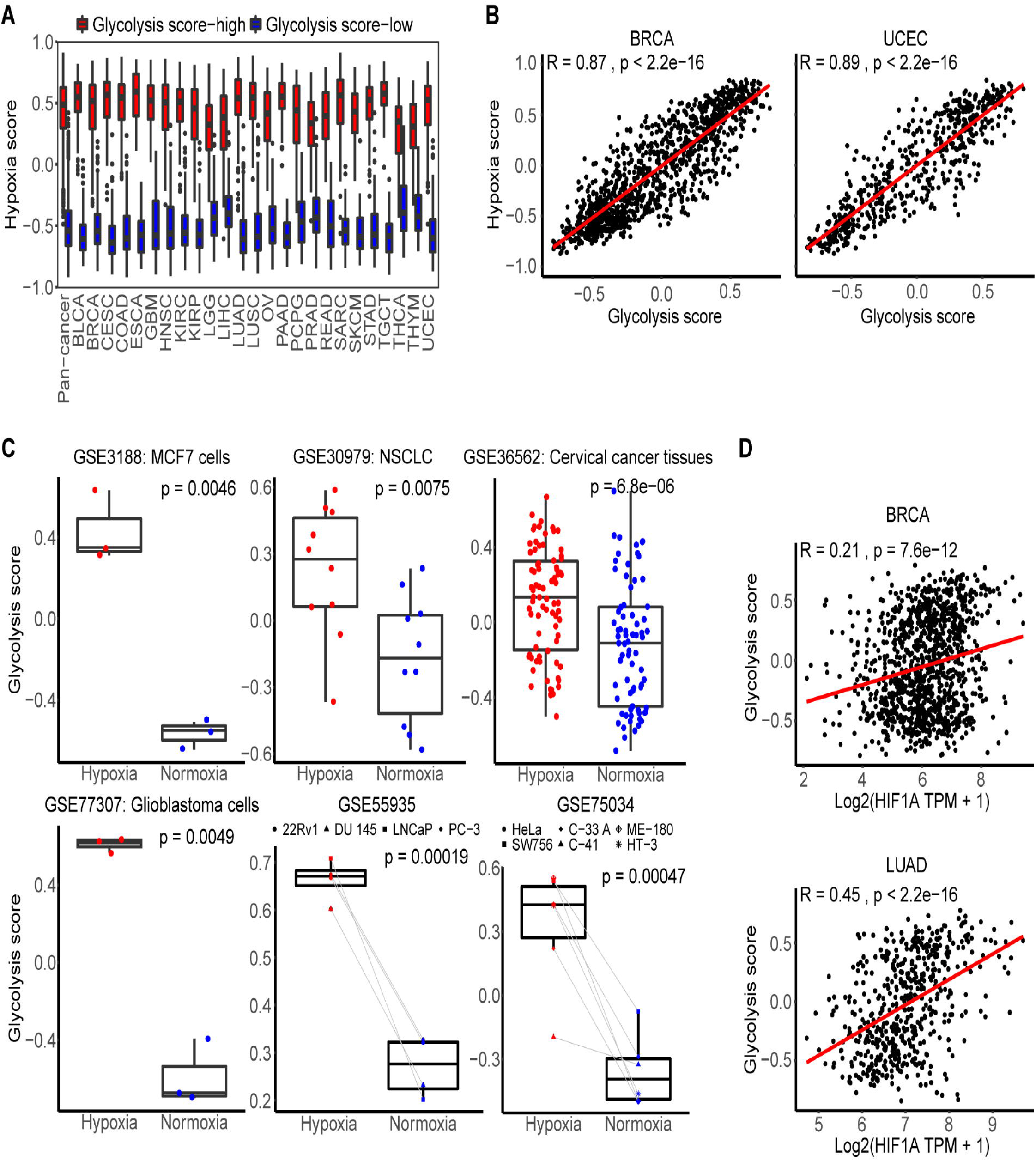
Association of glycolysis and hypoxia microenvironment. A. The distribution of hypoxia score in glycolysis high and low tumors across cancer types. B. Spearman correlation between tumor glycolysis score and hypoxia score in BRCA and UCEC. P < 0.05. C. The distribution of glycolysis score of cancer cell lines and tumor fragments under hypoxic and normoxic conditions in six datasets. Two-sided Student’s t-test and paired Student’s t-test were used to assess the difference. P□<□0.05. D. Spearman correlation between tumor glycolysis score and HIF1A expression in BRCA and LUAD. P < 0.05.

Further verification showed tumor cells and tissues under the hypoxic conditions harbored significantly higher glycolysis scores than those in normoxic state using six independent samples from GEO database (P< 0.01) (Fig.4C). Here, we also analyzed single HIF1A expression and the association with glycolysis score and found the positive correlation in 18 cancer types (r from 0.17 to 0.52, P < 0.05), such as r = 0.45 in LUAD (Fig.4D, Supplementary Fig.5C). All these results showed the strong correlations and associations between glycolysis and hypoxia.

### 2.5 Glycolysis-associated gene and pathway across cancer types

To address whether transcriptome features would differentiate between glycolysis high and low tumors, we compared the transcriptomes of the two tumor groups by analysis of differentially expressed genes (DEGs) (1.5-fold difference, adj.P < 0.05) coupled with reactome term enrichment analysis (adj.P < 0.01) of DEGs across cancer types (Supplementary Tab.5). From analysis of DEGs, we choose the genes that were co-upregulated and down-regulated genes in at least 13 cancer types. There were 251 genes up-regulated-in glycolysis high versus low tumors. In the pathway enrichment, not surprisingly, glycolysis/gluconeogenesis pathway as the most highly enriched pathway in our defined glycolysis high groups across cancers (Supplementary Tab.5). The matrix remodeling, cell cycle and gap junction pathways also showed significant enrichment in the glycolysis-high tumors (Fig.5A). Besides, DEGs that expressed higher in low glycolysis score tumors were enriched in ABC transporters in lipid homeostasis, biosynthesis of maresin like SPMs pathways (Supplementary Fig.6A).

**Fig.5:**
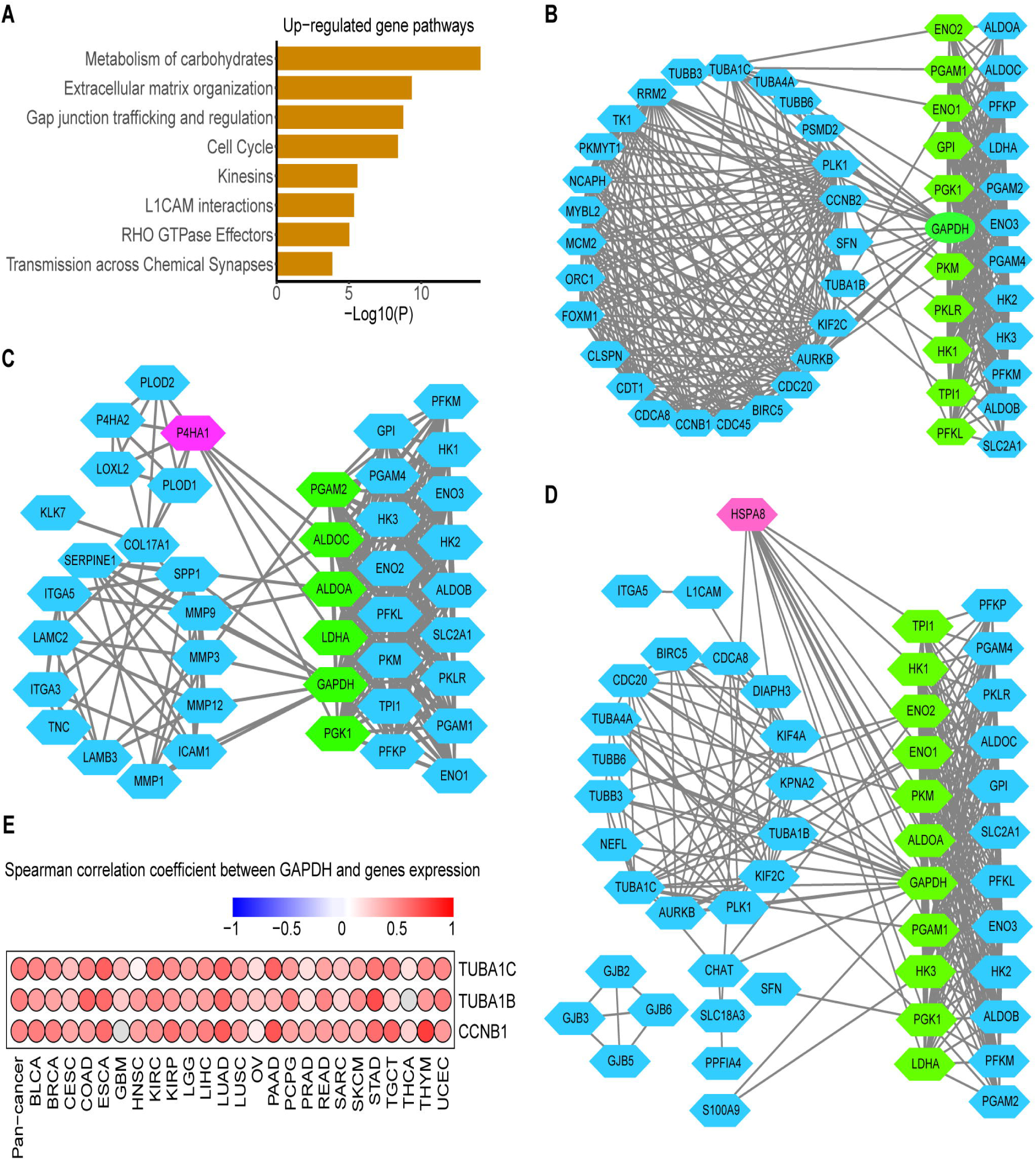
Glycolysis-associated mRNA and pathway signatures in hypoxia microenvironment. A. Reactome pathway enrichment of genes that were up-regulated in the glycolysis high tumors in at least 13 cancers. P < 10^−2^. B-D. Protein interaction network between genes in cell cycle (B), ECM remodeling (C) and GAP junking (D) with glycolysis genes. Blue nodes in left represent DEGs and blue nodes in right represent the glycolysis genes, green nodes represent key genes in glycolysis which mostly linked with DEGs and red nodes represent the genes with most linking with glycolysis genes. E. Spearman correlation between tumor GADPH and most glycolysis-related genes TUBA1C, TUBA1B and CCNB1 in cell cycle across cancer types.

### 2.6 Identification the glycolysis-associated genes

To identify genes most associated with glycolysis and explore interaction mechanism of these genes and glycolysis, we screened the genes according to the significance of their enrichment pathway. We regarded these up-regulated DEGs enriched in three most significantly (adj.P <10^−6^) enriched pathways (ECM remodeling, cell cycle and GAP junking) as candidate genes. The spearman correlation between above candidate genes with glycolysis score was carried out to screen the most glycolysis-related genes. TUBA1C (r was range from 0.29 to 0.66), TUBA1B (0.21-0.65), P4HA1 (0.25-0.74), HSPA8 (0.17-0.60), CCNB1 (0.22-0.62) as the top five genes were highly correlated with glycolysis score across cancers (Supplementary Fig.6B, Supplementary Tab.6).

To explore how glycolysis may influence these genes mRNA abundance or how glycolysis may be regulated by them. We constructed the protein interactions of candidate genes with glycolysis genes to confirm the protein relationship between them. As Figure 5 shows there were many linking among them, GADPH was identified the glycolysis genes to hold the most interaction with candidate genes while candidate gene HSPA8 and P4HA1 was interacted with most glycolysis genes (Fig.5B-D). As we know glycolysis enzyme could act nuclear function to regulate genes acting in cell cycle, such as GADPH^[31]^, so we carried out correlation analysis of mRNA expression between GAPDH and TUBA1C, TUBA1B, CCNB1 in cell cycle which are top genes were highly correlated with glycolysis score identified above and the results showed that GAPDH was positively correlated with them in 24∼25 cancer types, such as, correlation coefficient of GAPDH and CCNB1 as high as 0.77 in THYM, 0.68 in PAAD, 0.67 in LUAD (Fig.5E, P < 0.05).

### 2.7 Association of glycolysis and HSPA8, P4HA1

As HSPA8 and P4HA1, which are top genes were highly correlated with glycolysis score identified above, were also interacted with most glycolysis genes in the PPI analysis (Fig.5C-D), we hypothesized that HSPA8 and P4HA1 may be the potential factors influencing glycolysis. Thus, the correlation analysis of HSPA8 and P4HA1 with glycolysis genes was carried out. The results showed that P4HA1 was correlated with PGK1 (r = 0.16∼0.81) in 25 cancer types and correlated with LDHA (r = 0.13∼0.79) in 24 cancer types (P < 0.05) (Fig.6A). HSPA8 was associated with most of glycolysis genes like PGK1 (r = 0.21∼0.79) and PGAM1(r = 0.17∼0.80) across all cancer types (Fig.6B, P <□0.05).

**Fig.6:**
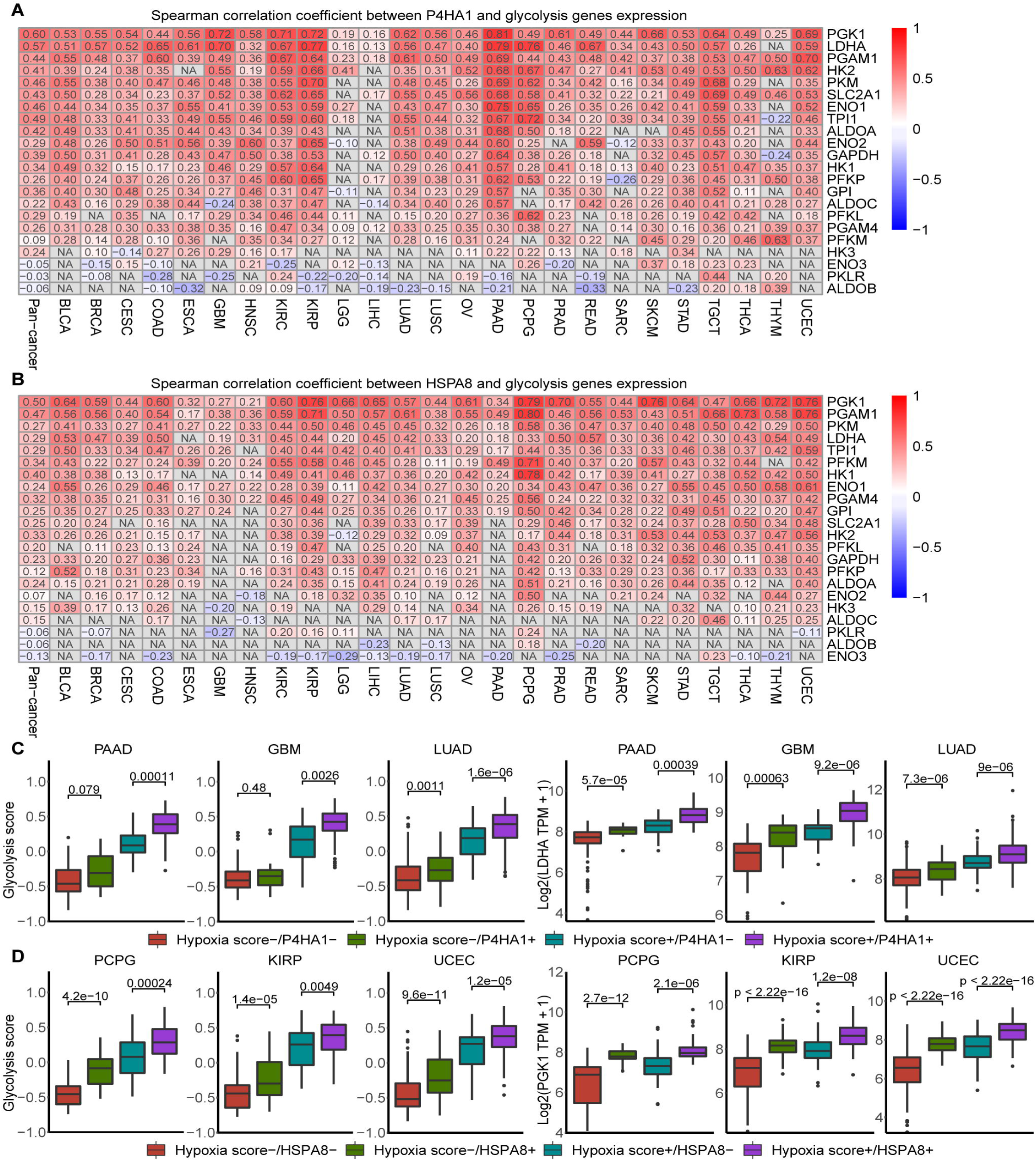
Association of glycolysis and P4HA1, HSPA8 in hypoxia microenvironment. A. Spearman correlation of between P4HA1 and 22 gene expression across cancer types. B. Spearman correlation of between HSPA8 and 22 gene expression across cancer types. C. Glycolysis score levels and LDHA mRNA abundance differ depending on hypoxia status and P4HA1 expression in three cancer types. Box plots represent the median (center line) and upper and lower quartiles (box limits). Hypoxia score+ indicates tumors with top 50% hypoxia score, Hypoxia score− indicates tumors with bottom 50% hypoxia score. P4HA1+ indicates tumors with top 50% P4HA1 expression in the Hypoxia score+ or Hypoxia score− groups, P4HA1− indicates tumors with bottom 50% P4HA1 expression in the Hypoxia score+ or Hypoxia score− groups. D. Glycolysis score levels and PGK1 mRNA abundance differ depending on hypoxia status and HSPA8 expression in three cancer types. Hypoxia score+ indicates tumors with top 50% hypoxia score, Hypoxia score− indicates tumors with bottom 50% hypoxia score. HSPA8+ indicates tumors with top 50% HSPA8 expression in the Hypoxia score+ or Hypoxia score− groups, HSPA8− indicates tumors with bottom 50% HSPA8 expression in the Hypoxia score+ or Hypoxia score− groups.

To further explore whether P4HA1 and HSPA8 affect glycolysis by regulation of glycolytic genes in hypoxia environment, we compared the differences of glycolytic score and key glycolysis gene expression at different hypoxia state and P4HA1 or HSPA8 expression patterns. As showed in Figure 6C, hypoxia with P4HA1 mRNA abundance significantly predicted glycolysis score and LDHA expression across cancer types, such as, GBM, LUAD and PAAD. The highest glycolysis score, LDHA and PGK1 mRNA abundance were observed under high hypoxia and high P4HA1 expression in some cancer types (Fig.6C, Supplementary Fig.6C, Supplementary Tab.7). The same distribution as P4HA1, HSPA8 was also proved to be a factor influencing glycolysis score and key glycolysis genes expression in different hypoxia state across cancer types. The highest glycolysis score, PGK1 and PGAM1 mRNA abundance were observed under high hypoxia and high HSPA8 expression across cancer types (Fig.6D, Supplementary Fig.6D, Supplementary Tab.7). To study the expression of HSPA8 and P4HA1 under hypoxia, the different expression of HSPA8 and P4HA1 was observed in cancer cell lines and tumor fragments of multiple cancer types under hypoxia compared with normoxic conditions from previous studies (GEO numbers see Methods). P4HA1 was up-regulated in cancer cells cultured with hypoxia across all single datasets (P <□0.05) (Supplementary Fig.7A), however, HSPA8 was showed the opposite trend in the same conditions in five datasets (P <□0.05) (Supplementary Fig.7B). Collectively, our data show that glycolysis activity are associated strongly with upregulated key genes HSPA8 and P4HA1, which are correlated with key glycolysis genes such as PGK1 and strongly associated with hypoxia state.

## 3. Discussion

The underlying biology of the Warburg effect has remained obscure, such as, the biological events controlled by the glycolysis pathway and the factors influencing glycolysis are not well defined. For example, although previous studies have linked glycolysis to hypoxia, we were completely blinded to their relationship in the large sample and data model, only existing biological knowledge was used for analysis individual glycolysis gene activated by hypoxia-inducible factor. For this, our bioinformatics approach uncovered the findings supporting the model that cancer cells favor glycolysis which is an important biological process that promote cancer cell proliferation in hypoxia condition. First, we interrogated the 22-gene expression representing glycolysis activity and verified our gene signature follow a remarkably robust performance in validation cohorts. Second, congruent with previous studies^[32]^, the high glycolysis tumors were associated with a markedly worse prognosis than the low glycolysis tumors in some cancer types. Third, cell proliferation signals and synthesis of nucleic acid and other macromolecules are active in glycolytic high tumors. Fourth, a strong positive correlation between the hypoxia signature and glycolysis activity is conserved in all cancer types. Fifth, we confirmed that GADPH may active the cell cycle by activation the transcription of cycle-dependent protein. Finally yet importantly, based on the strong correlation between HSPA8, P4HA1 and glycolysis score, glycolysis may be activated in hypoxic environment through other mechanisms not just HIF1A.

Our analysis shows that cell proliferation gene set is higher in glycolysis-high tumors relative to glycolysis-low tumors which is also mentioned by other studies^[33]^. As we know production of proteins, lipids and nucleic acids is essential for a successful replicative cell division thus promote cell proliferation^[34]^. It has been shown that the biosynthesis of these highly needed macromolecules is achieved mainly through acquisition and utilization of sources of nutrients from its metabolic intermediates of glycolysis^[34]^. Our results show that glycolysis-high tumors harbor more active of these cellular biosynthetic pathways than low tumors which can be partly explained the way of promoting cell proliferation by glycolysis. However, the mechanisms whereby tumor glycolysis induces tumor cell proliferation are incompletely understood. We hypothesize that in addition to provide the materials and energy needed for growth and division, tumor glycolysis may activate additional signaling networks that promote cell proliferation, such as, by increasing cell cycle-dependent enzyme activity.

The non-glycolytic functions of enzymes involved in glycolysis have recently been identified to be predominantly associated with the cancer development^[35, 36]^. CCNB1 is a regulatory protein involved in G2/M transition phase of the cell cycle and its encoding gene is highly correlated with glycolysis score (r = 0.22∼0.62) and GAPDH expression across tumors (r = 0.12∼0.77). Various non-glycolytic functions of GAPDH have been reported in cancer, such as, it has been reported that GAPDH overexpression in cell nucleus is associated with cell cycle via its effect on cyclin B-cdk1 activity^[31]^. Up-regulation of cell cycle related genes including CCNB1 is consistently associated with high-expressions of GAPDH in non-small-cell lung cancer^[37]^. Thus, combining these studies and high gene association supports the hypothesis that GAPDH may affect CCNB1 expression and cell cycle to promote cell proliferation. However, this study has been limited to measuring transcriptional levels based on the RNA-seq data, and further experimentation is required to verify this phenomenon including cellular protein levels.

To our surprise, we also find the enrichment of the TCA cycle and glutamic acid metabolism signatures are in glycolysis high tumors. Several previous studies comparing metabolic gene expression between tumors and normal tissues have identified suppression of OXPHOS as a recurrent metabolic phenotype in tumors^[38, 39]^. Although Warburg effect and these studies show glycolysis is a result of mitochondrial dysfunction, we now know that tumor glycolysis can proceed with functional cellular mitochondria and in fact may be an adaptive response for tumor survival^[40]^. It has also been known that most cancer cells do not have defects in mitochondrial metabolism^[41]^ and mitochondrial OXPHOS is also utilized by cancer cells^[42, 43]^. This observations indicate that a functional mitochondrial respiratory chain and a glutamine-derived carbon are required for proliferation of most cancer cells. Thus, this study indicates that mitochondrial OXPHOS is not mutually exclusive with glycolysis as routes for energy production in adapting to the tumor microenvironment.

Cancer cells may overcome growth factor dependence by deregulating inherent pathways that affect their metabolism, or by activating mutations in critical enzymes or transcription factor. Despite a wealth of data linking glycolysis with oncogenes and tumor suppressor genes^[44]^, most studies have focused on individual genes or in special cancer type. Thus, our study in a larger panel of tumors are to define the relationship between glycolysis activity and its influencing factor. In this study, we identify key modulators of glycolysis including the known factors and potential new ones uncovered by our analysis. It is well recognized that AKT and MYC are the most prevalent driving oncogenes in cancer and the potentially regulatory function affecting glycolysis^[45, 46] [47]^. In the present study, amplification in MYC and AKT1 show a strong consistency with transcript levels and glycolysis-high tumors harbor increased MYC and AKT1 expression. Besides, the expression of AKT1 and MYC are positively correlated with glycolysis score, suggesting that AKT1 and MYC activation may cooperated with each other and induce a maximally glycolytic state in some cancer type.

Although previous studies have linked glycolysis to hypoxia^[48]^, we were completely blinded to their relationship in the large sample model. Hypoxia is the result of an imbalance between oxygen delivery and oxygen consumption and is a common feature of solid tumors and is associated with their malignant phenotype^[49]^. This is the first study to identify glycolysis has such a strong correlation with hypoxia in a large data which indicates that glycolysis is the metabolic alteration responding to hypoxia in human cancers. However, most of this strong correlation may be contributed by the overlapped genes (LDHA, TPI1, ALDOA, ENO1, PGAM1 and SLC2A1) in hypoxia and glycolysis gene sets, which indicated these functional group of genes involved in glycolysis might consistently be regulated by hypoxia across cancer types. Perhaps the most important aspect of how these genes up-regulated in hypoxia is the activity of the HIF1 transcription factor^[50]^. HIF1 stimulates glycolytic energy production by transactivating genes involved in extracellular glucose import (such as SLC2A1^[51, 52]^) and can channel glucose into glycolysis by increasing the enzymes involved in this process (TPI1, HK2, PFK1, ALDOA, ENO1 and LDHA)^[20, 53]^. These previous researches suggest some genes involved in glycolysis can be directly regulated, at least in part, by HIF1, which can be partly explained the glycolysis genes are up-regulated in hypoxia condition and the high correlation between hypoxia and glycolysis. Our analysis of big transcriptomic profiles in a broader context clarifies that HIF1A expression is correlated with glycolysis score and activity of glycolysis is positively response to hypoxia in cancer cells from the tumor microenvironment. Based on previous studies, there are a portion of glycolytic genes not regulated by HIF1, we make assumptions that the glycolysis may be activated in hypoxic environment through other mechanisms not just HIF1A.

A comparison of gene expression in glycolysis high versus low groups revealed that the most significantly glycolysis-related gene HSPA8 and P4HA1 which are up-regulated in high glycolysis tumors. Given that HSPA8 is an chaperone by protection of the proteome from stress and the expression of HSPA8 is highly expressed in tumor tissue^[54]^, we assume it may play a critical role in mediating glycolysis coping with hypoxia pressure in tumor evolution. Our data reveals that high glycolysis score is associated with increased HSPA8 expression especially in hypoxia state which point to it as a potential mechanism that may be responsible for glycolysis. In addition, HSPA8 was up regulated in hypoxia-high across cancer types but down regulated in cancer cell cultured under hypoxia condition. This incongruent phenomenon in vitro and vivo experiment may be explained that cancer cells are coped with different pressure and duration in complex microenvironment in vitro culture or vivo tissues. We assume that HSPA8 may be a intermediate link responding to changing environmental pressures during the evolution of cancer and activating the glycolysis reaction, and futher study will be needed.

Previously, the increased P4HA1 expression correlates with poor prognosis has been demonstrated in several cancers, including breast cancer, oral squamous cell carcinoma, melanoma, and prostate cancer^[55-58]^. However, we know little about how P4HA1 promotes tumor progression in the past. A recent exciting study suggests that P4HA1 is essential for HIF-1 protein stability and is a new regulator of the HIF-1 pathway in breast cancer cells. They also show that overexpression P4HA1 in cancer cells induces LDHA mRNA levels, and P4HA1 expression correlates with LDHA mRNA levels in human breast cancer tissue^[59]^, which are consistent with our findings that highest LDHA mRNA abundance is observed under high P4HA1 expression condition and the expression of LDHA and P4HA1 are significantly correlated each other (r range from 0.13 to 0.79). P4HA1 is also induced upregulated expresssion in cancer cell cultured under hypoxia compared with normoxia condition. Thus, integrating previous studies with our new findings suggest that P4HA1 may be a new regulator responsible for cancer progression by upregulating the expression of key enzymes such as LDHA and inducing glycolysis in hypoxia environments. In summary, our findings identify P4HA1 and HSPA8 as the critical regulator in glycolysis and additional work should verify our results and characterize the role of them in hypoxia-response pathways to activate glycolysis.

## Conclusion

Overall, our study employs a novel biocomputational approach to define glycolytic activity by a gene set expression and offers a global picture of glycolysis in human cancer from the highly complex tumor microenvironment. This work shows that the hypoxia could activate glycolysis to regulate cell cycle and cell proliferation and suggests a combined therapeutic strategy, for upregulated glycolysis subtypes, tumors may be vulnerable to a combination therapy which targets the master regulatory factors of glycolysis in hypoxia. Ultimately, it will be necessary to validate our findings in large independent cohorts. For example, particularly those hypoxia-associated characteristics, will require long term, systematic in vitro modelling and will be the subject of future studies.

## 4. Materials and Methods

### 4.1 Multi-omics data and clinical data from TCGA and GEO

Molecular data, including mRNA expression, CNAs, SNAs and clinical data, including tumor stage and overall survival times, across 25 cancer types were downloaded from the TCGA data portal (https://portal.gdc.cancer.gov/). The GEO data was downloaded from NCBI GEO dataset (https://www.ncbi.nlm.nih.gov/) including GSE21217, GSE101644, GSE3188, GSE30979, GSE36562, GSE55935, GSE70051 and GSE75034 as validation data set.

### 4.2 Classification of glycolysis activity across different cancer types

According to the reactome pathway database, gene expression and their importance in cancer, we selected a 22-gene expression signature (SLC2A1, HK1, HK2, HK3, GPI, PFKL, PFKM, PFKP, ALDOA, ALDOB, ALDOC, TPI1, GAPDH, PGK1, PGAM1, PGAM4, ENO1, ENO2, ENO3, PKLR, PKM and LDHA) that belongs the glycolysis core pathway^[60]^.

To classify glycolysis status, we employed GSVA^[61]^ to calculate the glycolysis score in each cancer type based on the 22 mRNA-based glycolysis signatures (GSVA score was scaled from –1 to 1 in each sample). The top and bottom 30% score samples were assigned as glycolysis score-high and score-low groups in each cancer type, respectively. We included 25 cancer types with ≥ 30 samples in both glycolysis score-high and low groups for further analysis.

### 4.3 linical relevance analysis of glycolysis subtypes

We evaluated the associations of glycolysis score with two clinical features respectively: the patients’ overall survival time and tumor stage. The R package ‘‘survival’’ was used to perform the overall survival analysis and produce Kaplan-Meier survival plots. As for associations of glycolysis score and tumor stage analysis, T test was performed to access the glycolysis score in different tumor stage.

### 4.4 Genomic instability associated with glycolysis activity

The association of glycolysis activity with CNAs and SNVs was tested in BRCA, LUAD and UCEC with samples more than 500 by using all subjects who had both CNA and SNV data (n BRCA = 1091, n LUAD = 513, n UCEC = 543). CNA and SNV biases were assessed for all genes for which data were available within these tumor types. CNA biases were tested for 19729 genes in each tumor types. Copy-number changes were associated with glycolysis activity by a comparison of glycolysis scores between tumors that were copy-number neutral to those with a copy-number gain^[18]^ (or to those with a loss; T test). For each gene with SNV data, glycolysis scores were compared between tumors with an SNV and those without an SNV (T test). A Benjamini and Hochberg P-value adjustment was applied within each tumor-type cohort.

### 4.5 Pan-cancer associations of driver CNAs and glycolysis activity

Driver CNA associations were assessed within all 25 tumor types for previously described oncogenes and tumor-suppressor genes^[27]^. For each oncogene, differences in glycolysis scores were compared between tumors that were copy-number neutral and those with a copy-number gain. For each tumor-suppressor gene, differences in glycolysis scores were compared between tumors that were copy-number neutral and those with a copy-number loss (T test). The top 50 driver CNAs were plotted according to the number of tumor types in which each driver event had P < 0.05.

### 4.6 Gene set enrichment

To calculate single-sample gene set enrichment, we used the GSVA program to derive the absolute enrichment scores of gene sets from several publications and previously experimentally validated gene signatures from MsigDB(https://www.gsea-msigdb.org/gsea/msigdb/) as follows: tumor proliferation signature^[28]^, tumor inflammation signature^[62]^, cellular response to hypoxia, MYC targets, DNA replication, G2M checkpoint, P53 pathway, PI3K/AKT/mTOR pathway, IFNG signaling, genes up-regulated by ROS, DNA repair, degradation of ECM, collagen formation, angiogenesis, EMT markers, apoptosis, TGFB signaling, and metabolism related pathways.

To derive the GSVA score of each signature in each tumor sample, the normalized log_2_ (TPM+1) values were passed on as input for GSVA in the RNA-seq mode. Differentially enriched gene sets between the glycolysis high and low tumor groups were defined by GSVA adj.P < 0.05 (we used limma R pachage because the GSVA scores were normally distributed around zero)^[63]^.

### 4.7 Spearman correlation of glycolysis score and hypoxia score

We selected a 14-gene expression signature (ALDOA, MIF, TUBB6, P4HA1, SLC2A1, PGAM1, ENO1, LDHA, CDKN3, TPI1, NDRG1, VEGFA, ACOT7 and ADM)^[28, 29]^ that has been shown to perform the best when classifying hypoxia status. The GSVA was also employed to calculate the hypoxia score in each cancer type based on the gene signatures. The spearman correlation was calculated between glycolysis score and hypoxia score across cancer types (P < 0.05).

### 4.8 Identification of genes and pathways alterations between glycolysis score-high and low tumors

The difference of gene expression between glycolysis high and low groups was analyzed by edgeR (1.5-fold difference, adjust P < 0.05). We choose the genes co-upregulated in at least 13 cancer types as glycolysis positively related genes. Reactome pathway database (https://reactome.org/) is an integrated database containing advanced functional information for the systematic analysis of gene functions, biological pathways and other research. Metascape (http://metascape.org) is an online resource to perform the gene annotation and functional enrichment analysis. We used Metascape to perform the pathway enrichment with the threshold of P value□<□0.01 and the number of enriched genes□≥ □3 were concerned as significant.

## Acknowledgements

This work was supported by the National Key R&D Program of China (2018YFC0910201), the Key R&D Program of Guangdong Province (2019B020226001), and the Science and the Technology Planning Project of Guangzhou (201704020176).

## Author contributions

J.W., K.H., and H.D. designed the study. J.W. and K.H. conducted the study. Y.B. and Z.C. collected data. K.H., M.H. and S.L. interpreted the data. J.W. and M.H. wrote the manuscript with the support of H.D. take responsibility for the integrity of the data analysis.

## Supplementary information

### Supplementary Figures

Supplementary Fig.1: 22-gene signature for glycolysis activity and clinical significance of glycolysis.

A. Samples are ordered from lowest to highest glycolysis score with 22-gene expression distribution in pan-cancer. B. Association of glycolysis score with patient clinical stages across cancer types. T-test was used to assess the difference (P□<□0.05).

Supplementary Fig.2: Characteristics of CNAs and TMB associated with glycolysis activity.

A. Differential expression of MYC and AKT1 between amplification and neutral tumors across cancer types. B. Spearman correlation between MYC, AKT1 expression with glycolysis score across cancer types. C. High glycolysis tumors harbor with high TMB in some cancer types (P < 0.05).

Supplementary Fig.3: The spearman correlation between tumor proliferation signatures and glycolysis score across cancer types.

Supplementary Fig.4: The spearman correlation between hypoxia score and glycolysis score across cancer types.

Supplementary Fig 5: Association between glycolysis and hypoxia across cancer types.

A. Spearman correlation between glycolysis gene and hypoxia score. B. Spearman correlation between hypoxia gene and glycolysis score. C. Spearman correlation between HIF1A and glycolysis score.

Supplementary Fig 6: Glycolysis-associated transcriptome signatures.

A. Pathways down-regulated in glycolysis high tumors. B. Spearman correlation between glycolysis score and DEGs (enrich in ECM remodeling, cell cycle and GAP junking).

C. PGK1 mRNA levels differ depending on hypoxia and PAHA1 expression. D. PGAM1 mRNA levels differ depending on hypoxia and HSPA8 expression.

Supplementary Fig 7: Expression pattern of PAHA1 and HSPA8 in hypoxic and normoxic conditions based on previous study (see Method). A. Expression of PAHA1 in hypoxia and normoxic conditions. B. Expression of HSPA8 in hypoxia and normoxic conditions.

### Supplementary Tables

Supplementary Table 1: TCGA tumor type descriptions and the sample size.

Supplementary Table 2: Glycolysis-associated CNAs and SNAs.

Supplementary Table 3: Pan-cancer regions of significant CNAs.

Supplementary Table 4: Signatures of cancer hallmarks.

Supplementary Table 5: DEGs and significantly enriched pathways in glycolysis-high versus glycolysis-low tumors.

Supplementary Table 6: Spearman correlation between DEGs with glycolysis score across cancer types.

Supplementary Table 7: Association of glycolysis and P4HA1, HSPA8 in hypoxia microenvironment.

## References

[1] Pavlova NN, Thompson CB. The Emerging Hallmarks of Cancer Metabolism. Cell Metab. 2016. 23(1): 27–47.

[2] Hanahan D, Weinberg RA. Hallmarks of cancer: the next generation. Cell. 2011. 144(5): 646–74.

[3] Vander Heiden MG, Cantley LC, Thompson CB. Understanding the Warburg effect: the metabolic requirements of cell proliferation. Science. 2009. 324(5930): 1029–33.

[4] Li Z, Zhang H. Reprogramming of glucose, fatty acid and amino acid metabolism for cancer progression. Cell Mol Life Sci. 2016. 73(2): 377–92.

[5] Cascone T, McKenzie JA, Mbofung RM, et al. Increased Tumor Glycolysis Characterizes Immune Resistance to Adoptive T Cell Therapy. Cell Metab. 2018. 27(5): 977–987.e4.

[6] Ruprecht B, Zaal EA, Zecha J, et al. Lapatinib Resistance in Breast Cancer Cells Is Accompanied by Phosphorylation-Mediated Reprogramming of Glycolysis. Cancer Res. 2017. 77(8): 1842–1853.

[7] Fonti R, Pellegrino S, Catalano L, Pane F, Del Vecchio S, Pace L. Visual and volumetric parameters by 18F-FDG-PET/CT: a head to head comparison for the prediction of outcome in patients with multiple myeloma. Ann Hematol. 2020. 99(1): 127–135.

[8] Wu LL, Liang JH, Wang L, Xu W, Ding CY. [Prognostic value of pretreatment (18)F-FDG PET-CT metabolic parameters in patients with advanced extranodal NK/T cell lymphoma]. Zhonghua Zhong Liu Za Zhi. 2019. 41(11): 831–836.

[9] Salazar-Roa M, Malumbres M. Fueling the Cell Division Cycle. Trends Cell Biol. 2017. 27(1): 69–81.

[10] Sun S, Li H, Chen J, Qian Q. Lactic Acid: No Longer an Inert and End-Product of Glycolysis. Physiology (Bethesda). 2017. 32(6): 453–463.

[11] Corbet C, Feron O. Tumour acidosis: from the passenger to the driver’s seat. Nat Rev Cancer. 2017. 17(10): 577–593.

[12] Eriksson M, Ambroise G, Ouchida AT, et al. Effect of Mutant p53 Proteins on Glycolysis and Mitochondrial Metabolism. Mol Cell Biol. 2017. 37(24).

[13] Tateishi K, Iafrate AJ, Ho Q, et al. Myc-Driven Glycolysis Is a Therapeutic Target in Glioblastoma. Clin Cancer Res. 2016. 22(17): 4452–65.

[14] Xie Y, Shi X, Sheng K, et al. PI3K/Akt signaling transduction pathway, erythropoiesis and glycolysis in hypoxia (Review). Mol Med Rep. 2019. 19(2): 783–791.

[15] Chang CH, Qiu J, O’Sullivan D, et al. Metabolic Competition in the Tumor Microenvironment Is a Driver of Cancer Progression. Cell. 2015. 162(6): 1229–41.

[16] Fischer GM, Vashisht Gopal YN, McQuade JL, Peng W, DeBerardinis RJ, Davies MA. Metabolic strategies of melanoma cells: Mechanisms, interactions with the tumor microenvironment, and therapeutic implications. Pigment Cell Melanoma Res. 2018. 31(1): 11–30.

[17] Black JC, Atabakhsh E, Kim J, et al. Hypoxia drives transient site-specific copy gain and drug-resistant gene expression. Genes Dev. 2015. 29(10): 1018–31.

[18] Bhandari V, Hoey C, Liu LY, et al. Molecular landmarks of tumor hypoxia across cancer types. Nat Genet. 2019. 51(2): 308–318.

[19] Schito L, Rey S. Cell-Autonomous Metabolic Reprogramming in Hypoxia. Trends Cell Biol. 2018. 28(2): 128–142.

[20] Denko NC. Hypoxia, HIF1 and glucose metabolism in the solid tumour. Nat Rev Cancer. 2008. 8(9): 705–13.

[21] Maher JC, Wangpaichitr M, Savaraj N, Kurtoglu M, Lampidis TJ. Hypoxia-inducible factor-1 confers resistance to the glycolytic inhibitor 2-deoxy-D-glucose. Mol Cancer Ther. 2007. 6(2): 732–41.

[22] Flaig TW, Glodé M, Gustafson D, et al. A study of high-dose oral silybin-phytosome followed by prostatectomy in patients with localized prostate cancer. Prostate. 2010. 70(8): 848–55.

[23] Heiden BT, Chen G, Hermann M, et al. 18F-FDG PET intensity correlates with a hypoxic gene signature and other oncogenic abnormalities in operable non-small cell lung cancer. PLoS One. 2018. 13(7): e0199970.

[24] Vlassenko AG, McConathy J, Couture LE, et al. Aerobic Glycolysis as a Marker of Tumor Aggressiveness: Preliminary Data in High Grade Human Brain Tumors. Dis Markers. 2015. 2015: 874904.

[25] Heiden BT, Patel N, Nancarrow DJ, et al. Positron Emission Tomography 18F-Fluorodeoxyglucose Uptake Correlates with KRAS and EMT Gene Signatures in Operable Esophageal Adenocarcinoma. J Surg Res. 2018. 232: 621–628.

[26] Jadvar H, Alavi A, Gambhir SS. 18F-FDG uptake in lung, breast, and colon cancers: molecular biology correlates and disease characterization. J Nucl Med. 2009. 50(11): 1820–7.

[27] Zack TI, Schumacher SE, Carter SL, et al. Pan-cancer patterns of somatic copy number alteration. Nat Genet. 2013. 45(10): 1134–40.

[28] Thienpont B, Steinbacher J, Zhao H, et al. Tumour hypoxia causes DNA hypermethylation by reducing TET activity. Nature. 2016. 537(7618): 63–68.

[29] Buffa FM, Harris AL, West CM, Miller CJ. Large meta-analysis of multiple cancers reveals a common, compact and highly prognostic hypoxia metagene. Br J Cancer. 2010. 102(2): 428–35.

[30] Ye Y, Hu Q, Chen H, et al. Characterization of Hypoxia-associated Molecular Features to Aid Hypoxia-Targeted Therapy. Nat Metab. 2019. 1(4): 431–444.

[31] Carujo S, Estanyol JM, Ejarque A, Agell N, Bachs O, Pujol MJ. Glyceraldehyde 3-phosphate dehydrogenase is a SET-binding protein and regulates cyclin B-cdk1 activity. Oncogene. 2006. 25(29): 4033–42.

[32] Riester M, Xu Q, Moreira A, Zheng J, Michor F, Downey RJ. The Warburg effect: persistence of stem-cell metabolism in cancers as a failure of differentiation. Ann Oncol. 2018. 29(1): 264–270.

[33] Xiao H, Wang J, Yan W, et al. GLUT1 regulates cell glycolysis and proliferation in prostate cancer. Prostate. 2018. 78(2): 86–94.

[34] Zhu J, Thompson CB. Metabolic regulation of cell growth and proliferation. Nat Rev Mol Cell Biol. 2019. 20(7): 436–450.

[35] Chang YC, Yang YC, Tien CP, Yang CJ, Hsiao M. Roles of Aldolase Family Genes in Human Cancers and Diseases. Trends Endocrinol Metab. 2018. 29(8): 549–559.

[36] Hu JW, Sun P, Zhang DX, Xiong WJ, Mi J. Hexokinase 2 regulates G1/S checkpoint through CDK2 in cancer-associated fibroblasts. Cell Signal. 2014. 26(10): 2210–6.

[37] Wang D, Moothart DR, Lowy DR, Qian X. The expression of glyceraldehyde-3-phosphate dehydrogenase associated cell cycle (GACC) genes correlates with cancer stage and poor survival in patients with solid tumors. PLoS One. 2013. 8(4): e61262.

[38] Reznik E, Wang Q, La K, Schultz N, Sander C. Mitochondrial respiratory gene expression is suppressed in many cancers. Elife. 2017. 6.

[39] Hakimi AA, Reznik E, Lee CH, et al. An Integrated Metabolic Atlas of Clear Cell Renal Cell Carcinoma. Cancer Cell. 2016. 29(1): 104–116.

[40] Frezza C, Gottlieb E. Mitochondria in cancer: not just innocent bystanders. Semin Cancer Biol. 2009. 19(1): 4–11.

[41] Koppenol WH, Bounds PL, Dang CV. Otto Warburg’s contributions to current concepts of cancer metabolism. Nat Rev Cancer. 2011. 11(5): 325–37.

[42] LeBleu VS, O’Connell JT, Gonzalez Herrera KN, et al. PGC-1α mediates mitochondrial biogenesis and oxidative phosphorylation in cancer cells to promote metastasis. Nat Cell Biol. 2014. 16(10): 992–1003, 1-15.

[43] Viale A, Corti D, Draetta GF. Tumors and mitochondrial respiration: a neglected connection. Cancer Res. 2015. 75(18): 3685–6.

[44] Kroemer G, Pouyssegur J. Tumor cell metabolism: cancer’s Achilles’ heel. Cancer Cell. 2008. 13(6): 472–82.

[45] Stine ZE, Walton ZE, Altman BJ, Hsieh AL, Dang CV. MYC, Metabolism, and Cancer. Cancer Discov. 2015. 5(10): 1024–39.

[46] Wong K, Liao JZ, Verheyen EM. A positive feedback loop between Myc and aerobic glycolysis sustains tumor growth in a Drosophila tumor model. Elife. 2019. 8.

[47] Wieman HL, Wofford JA, Rathmell JC. Cytokine stimulation promotes glucose uptake via phosphatidylinositol-3 kinase/Akt regulation of Glut1 activity and trafficking. Mol Biol Cell. 2007. 18(4): 1437–46.

[48] Haider S, McIntyre A, van Stiphout RG, et al. Genomic alterations underlie a pan-cancer metabolic shift associated with tumour hypoxia. Genome Biol. 2016. 17(1): 140.

[49] Brizel DM, Rosner GL, Prosnitz LR, Dewhirst MW. Patterns and variability of tumor oxygenation in human soft tissue sarcomas, cervical carcinomas, and lymph node metastases. Int J Radiat Oncol Biol Phys. 1995. 32(4): 1121–5.

[50] Iyer NV, Kotch LE, Agani F, et al. Cellular and developmental control of O2 homeostasis by hypoxia-inducible factor 1 alpha. Genes Dev. 1998. 12(2): 149–62.

[51] Maxwell PH, Dachs GU, Gleadle JM, et al. Hypoxia-inducible factor-1 modulates gene expression in solid tumors and influences both angiogenesis and tumor growth. Proc Natl Acad Sci U S A. 1997. 94(15): 8104–9.

[52] Chen C, Pore N, Behrooz A, Ismail-Beigi F, Maity A. Regulation of glut1 mRNA by hypoxia-inducible factor-1. Interaction between H-ras and hypoxia. J Biol Chem. 2001. 276(12): 9519–25.

[53] Semenza GL, Roth PH, Fang HM, Wang GL. Transcriptional regulation of genes encoding glycolytic enzymes by hypoxia-inducible factor 1. J Biol Chem. 1994. 269(38): 23757–63.

[54] Shan N, Zhou W, Zhang S, Zhang Y. Identification of HSPA8 as a candidate biomarker for endometrial carcinoma by using iTRAQ-based proteomic analysis. Onco Targets Ther. 2016. 9: 2169–79.

[55] Gilkes DM, Chaturvedi P, Bajpai S, et al. Collagen prolyl hydroxylases are essential for breast cancer metastasis. Cancer Res. 2013. 73(11): 3285–96.

[56] Xiong G, Deng L, Zhu J, Rychahou PG, Xu R. Prolyl-4-hydroxylase α subunit 2 promotes breast cancer progression and metastasis by regulating collagen deposition. BMC Cancer. 2014. 14: 1.

[57] Atkinson A, Renziehausen A, Wang H, et al. Collagen Prolyl Hydroxylases Are Bifunctional Growth Regulators in Melanoma. J Invest Dermatol. 2019. 139(5): 1118–1126.

[58] Kappler M, Kotrba J, Kaune T, et al. P4HA1: A single-gene surrogate of hypoxia signatures in oral squamous cell carcinoma patients. Clin Transl Radiat Oncol. 2017. 5: 6–11.

[59] Xiong G, Stewart RL, Chen J, et al. Collagen prolyl 4-hydroxylase 1 is essential for HIF-1α stabilization and TNBC chemoresistance. Nat Commun. 2018. 9(1): 4456.

[60] Fabregat A, Sidiropoulos K, Viteri G, et al. Reactome pathway analysis: a high-performance in-memory approach. BMC Bioinformatics. 2017. 18(1): 142.

[61] Hänzelmann S, Castelo R, Guinney J. GSVA: gene set variation analysis for microarray and RNA-seq data. BMC Bioinformatics. 2013. 14: 7.

[62] Danaher P, Warren S, Lu R, et al. Pan-cancer adaptive immune resistance as defined by the Tumor Inflammation Signature (TIS): results from The Cancer Genome Atlas (TCGA). J Immunother Cancer. 2018. 6(1): 63.

[63] Hugo W, Zaretsky JM, Sun L, et al. Genomic and Transcriptomic Features of Response to Anti-PD-1 Therapy in Metastatic Melanoma. Cell. 2017. 168(3): 542.

