## Supplementary figures and images for "Persistent cell proliferation signals correlates with increased glycolysis in tumor hypoxia microenvironment across cancer types"

### SF1.tif

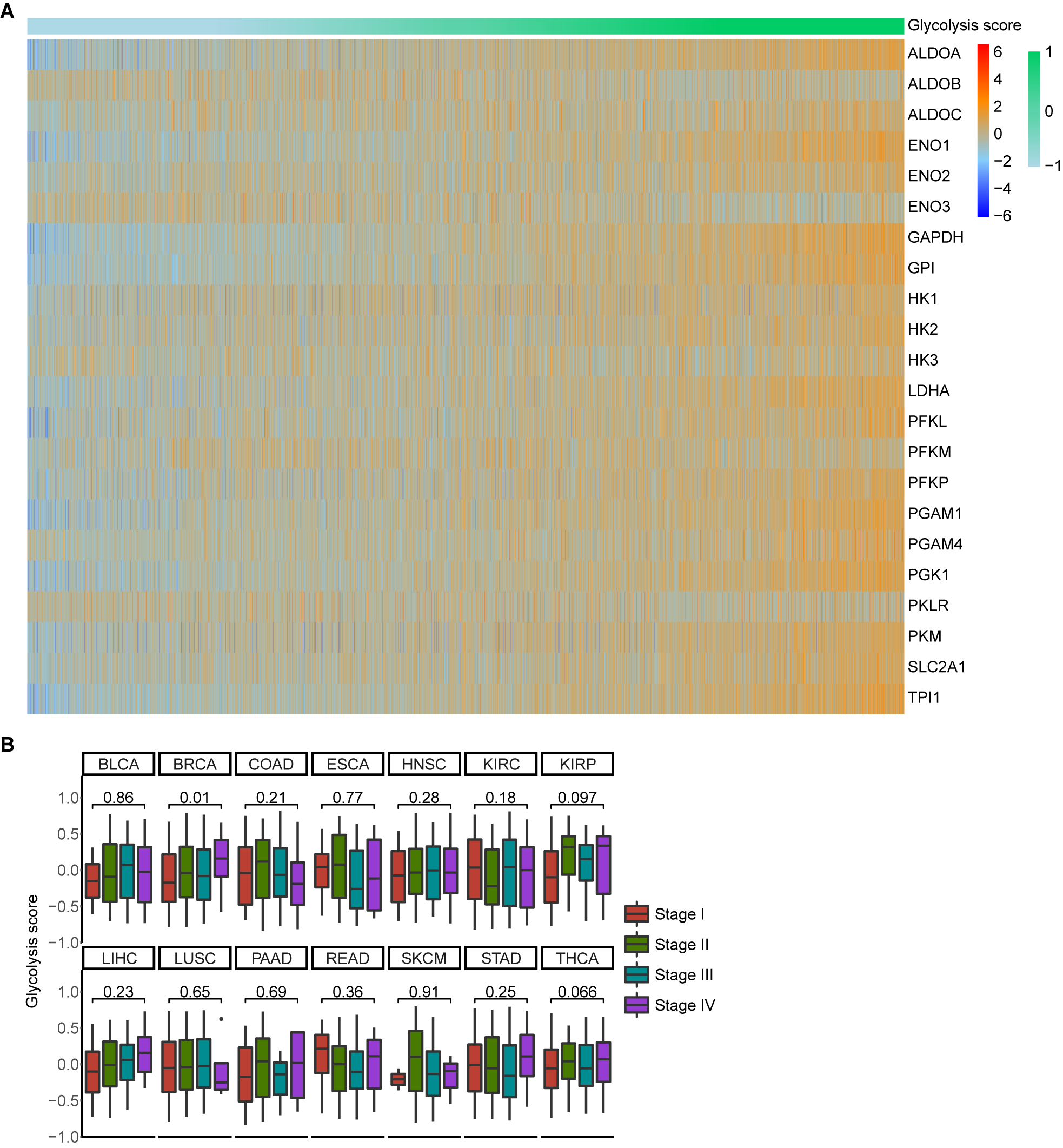

### SF2.tif

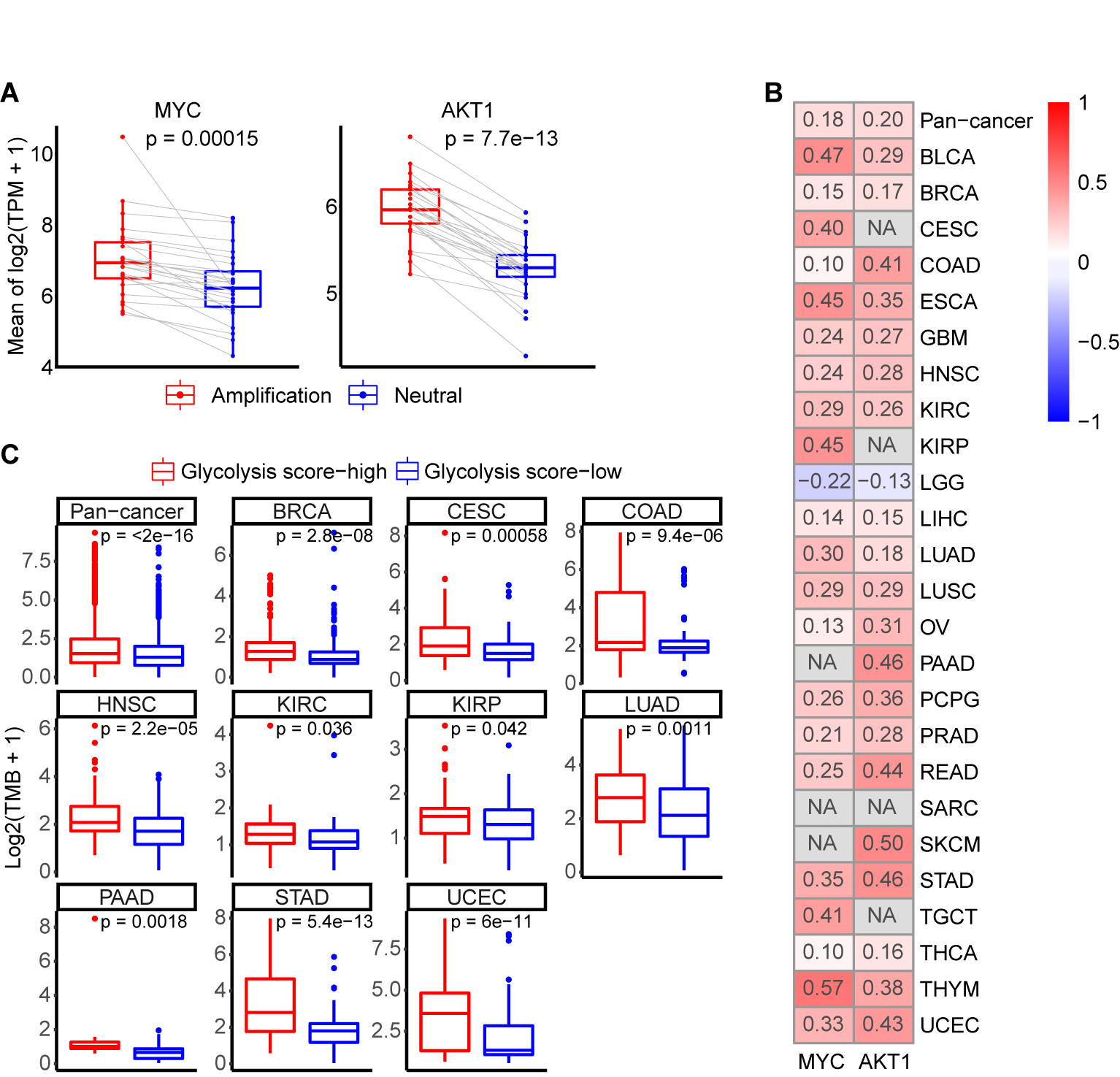

### SF3.tif

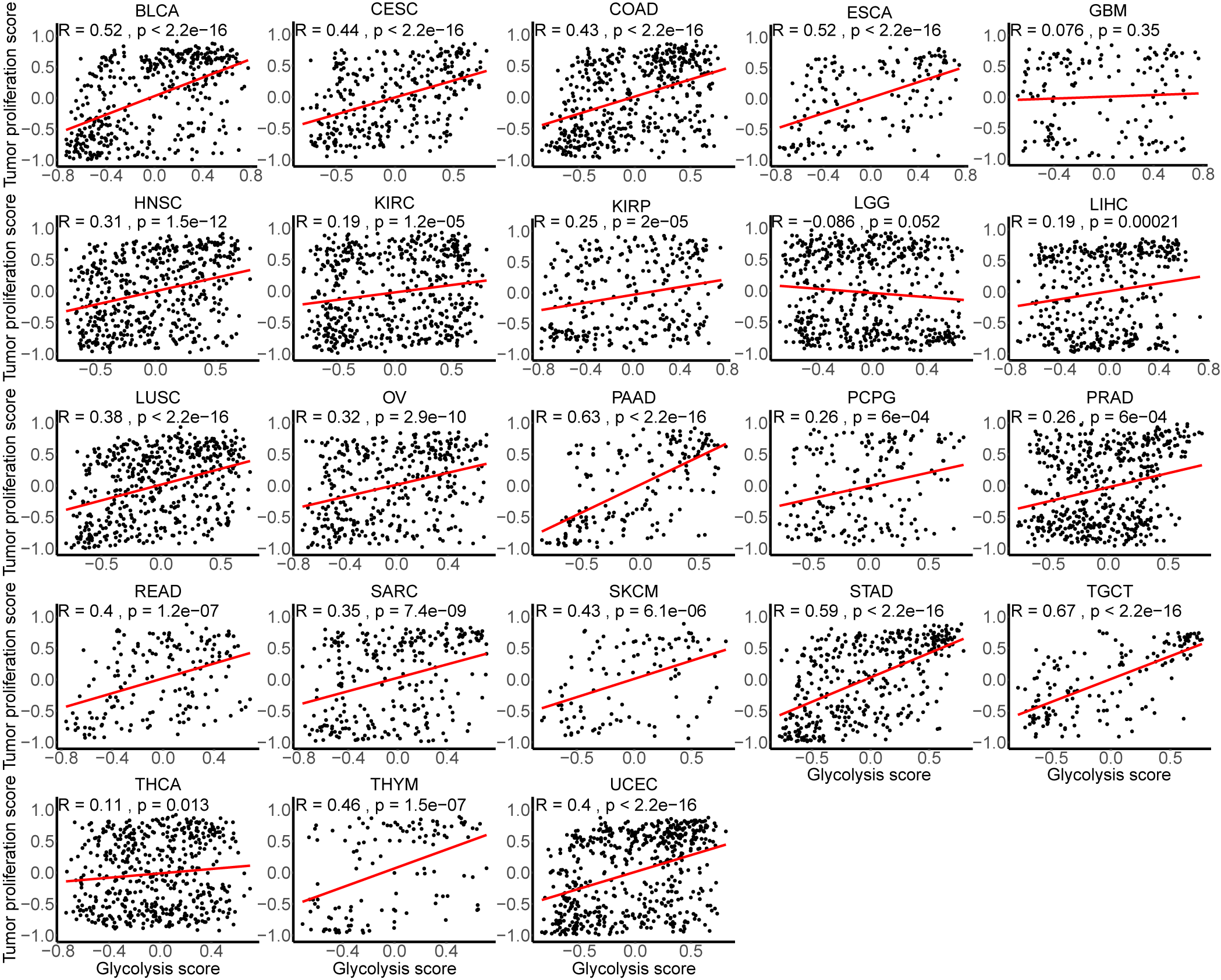

### SF4.tif

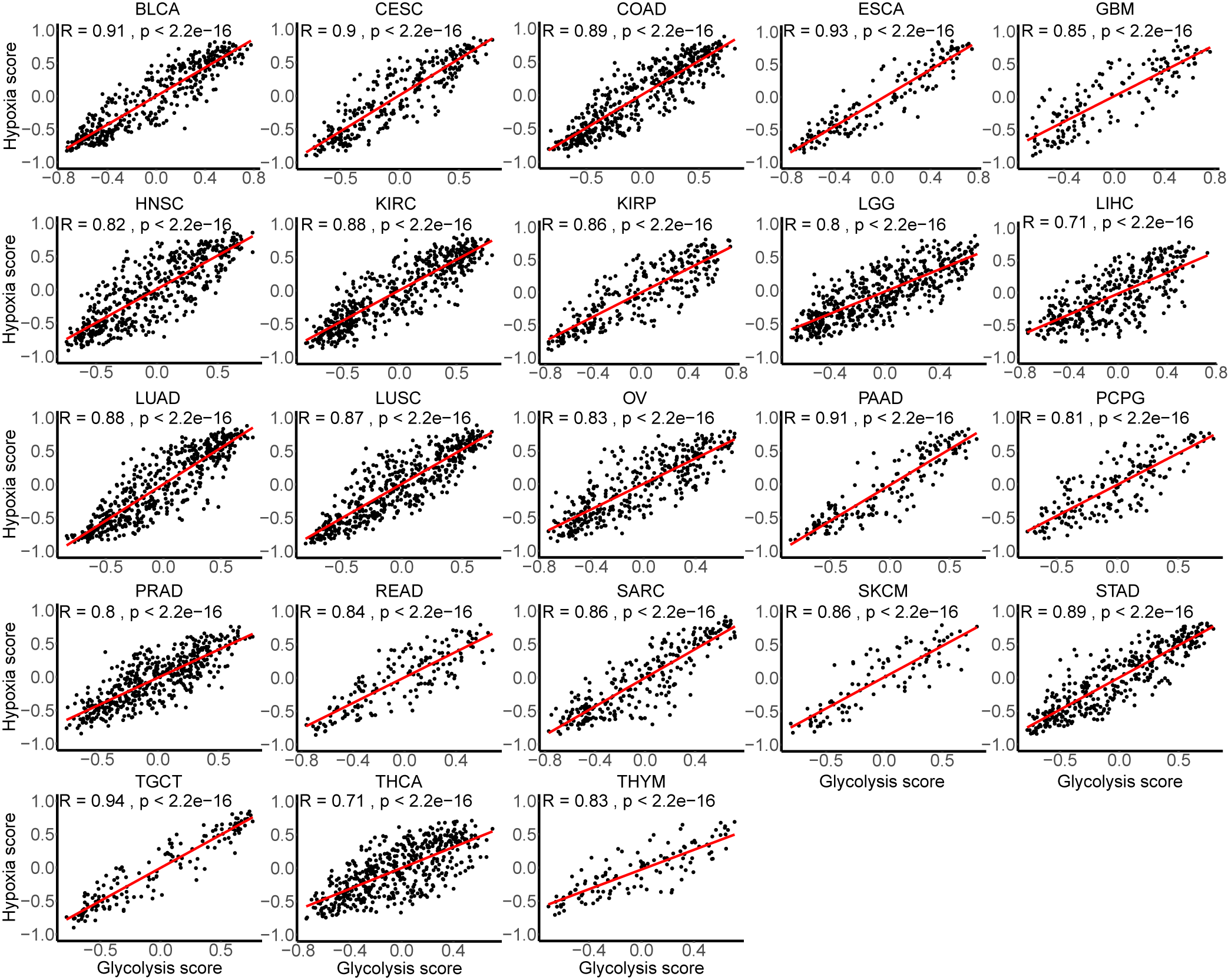

### SF5.tif

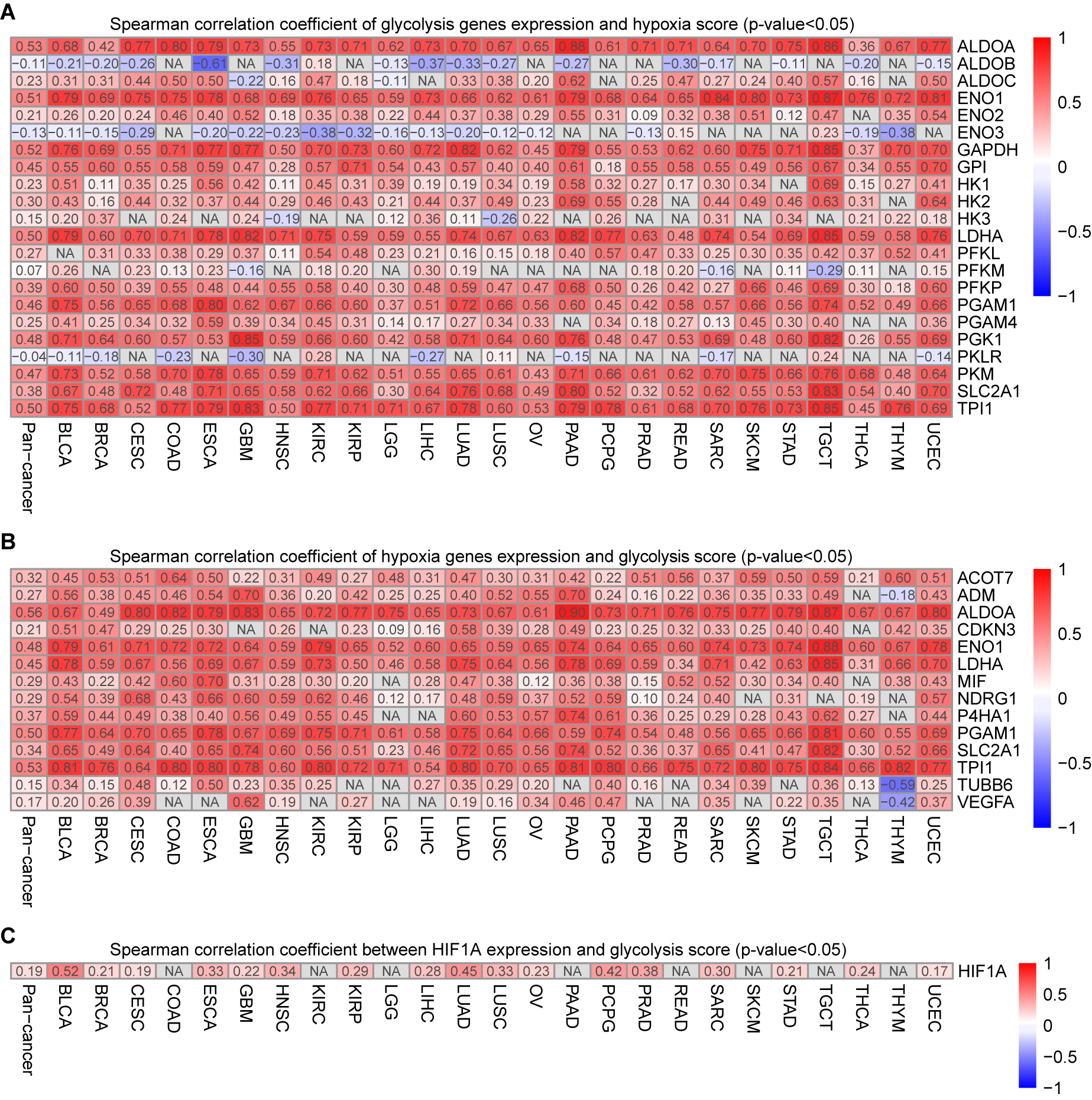

### SF6.tif

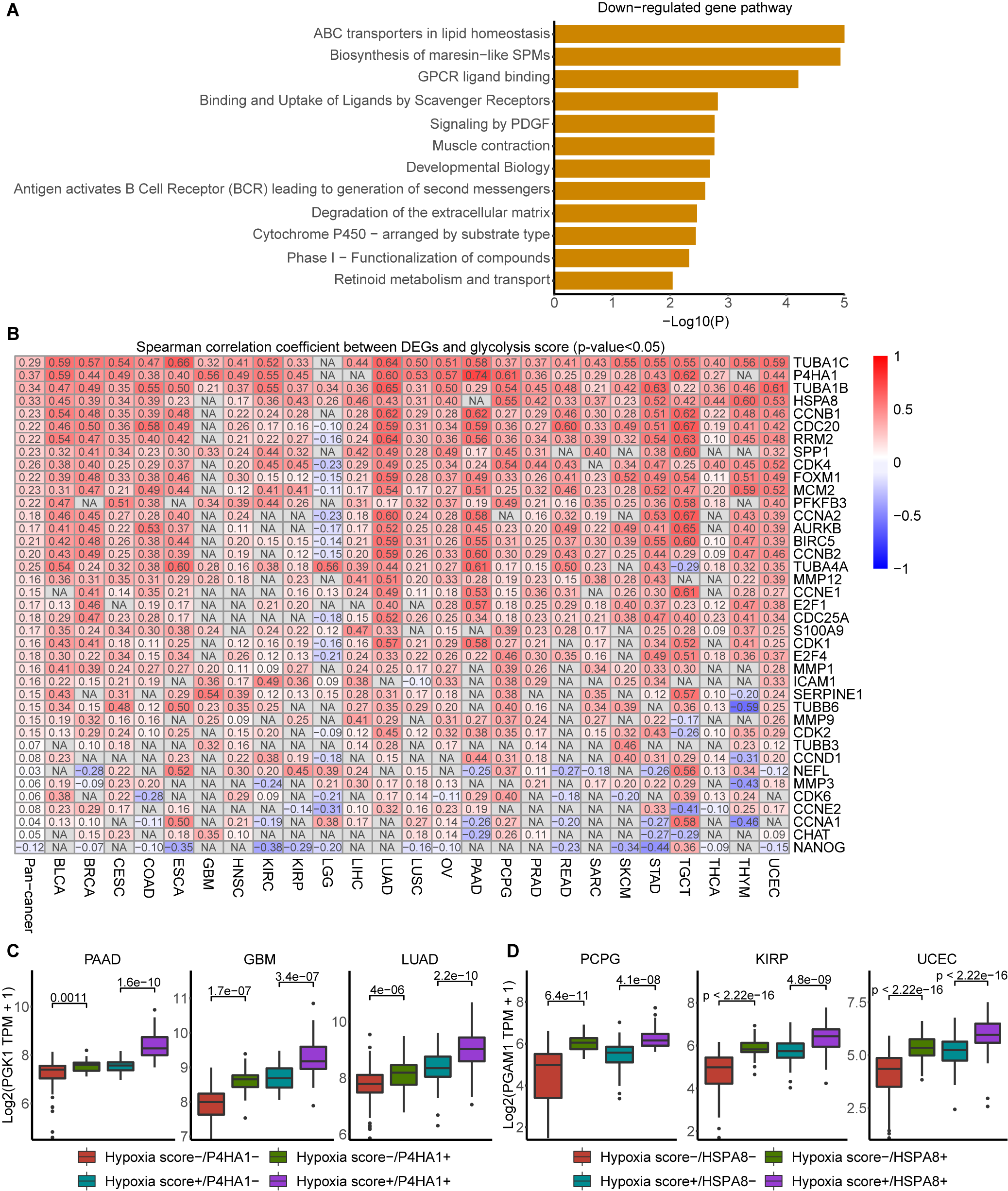

### SF7.tif

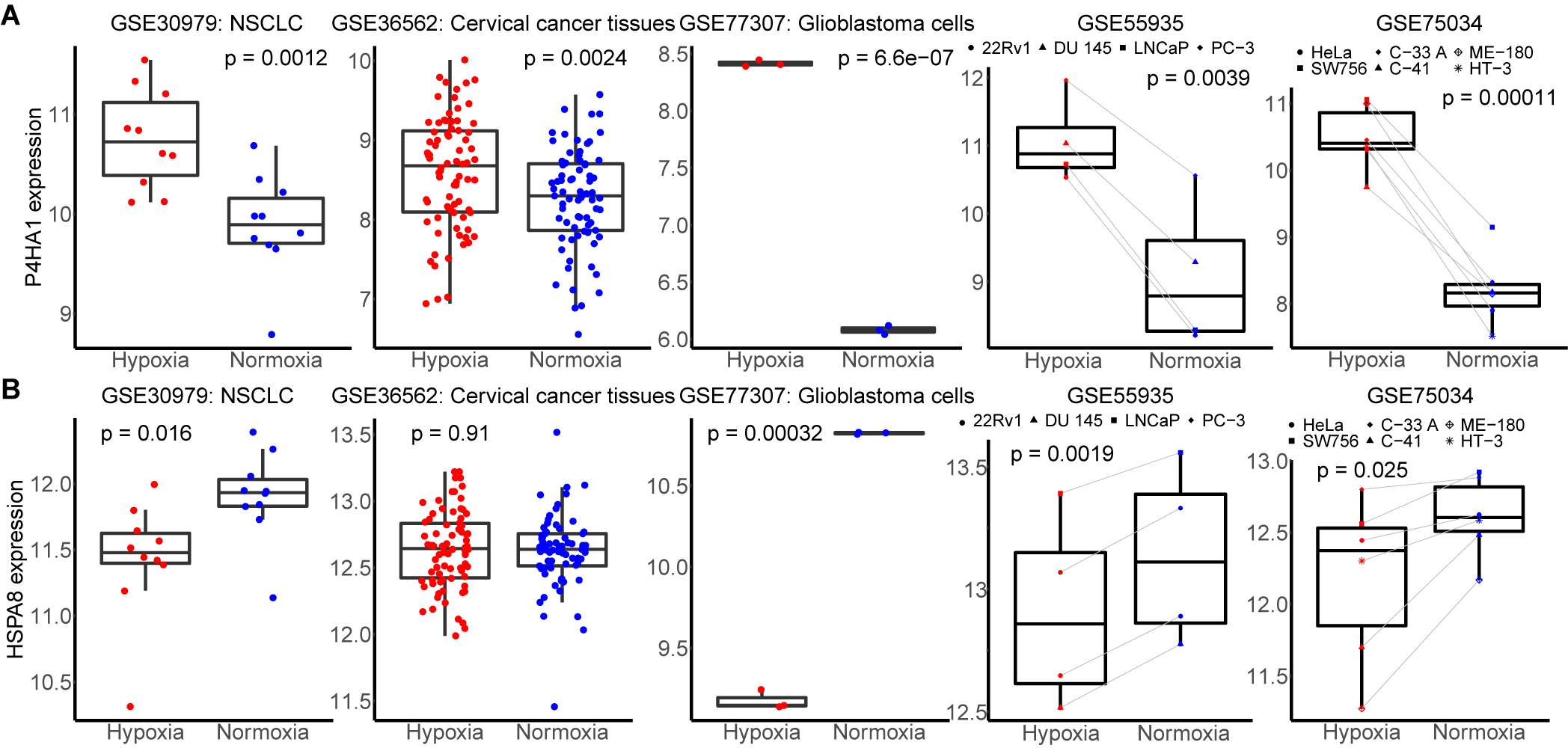
